# Timing and Delivery route effects of Cecal Microbiome transplants on *Salmonella* Typhimurium infections in Chickens

**DOI:** 10.1101/2022.08.08.502921

**Authors:** Sian Pottenger, Amyleigh Watts, Amy Wedley, Sue Jopson, Alistair C Darby, Paul Wigley

## Abstract

**Background:** Exposure to microbes early in life has long-lasting effects on microbial community structure and function of the microbiome. However, in commercial poultry settings chicks are reared as a single-age cohort with no exposure to adult birds which can have profound effects on microbiota development and subsequent pathogen challenge. Microbiota manipulation is a proven and promising strategy to help reduce pathogen load and transmission within broiler flocks.

**Results:** Manipulation of the microbiota between 4 and 72 hours of hatch markedly reduces faecal shedding and colonisation with the foodborne pathogen *Salmonella enterica* serovar Typhimurium (ST4/74). Administration route has minimal effect on the protection conferred with fewer birds in transplant groups shown to shed ST4/74 in the faeces compared to PBS-gavaged control birds. Analysis of the microbiome following transplantation demonstrated that the relative abundance of the anti-inflammatory bacterium *Faecalibacterium prausnitzii* was significantly higher in CMT groups compared to PBS controls. The presence of *F. prausnitzii* was also shown to increase in PBS-challenged birds compared to unchallenged birds potentially indicating a role of this bacterium in limiting *Salmonella* infections.

**Conclusions:** This study highlights the efficacy of using microbiome transplants as a means to reduce colonisation and shedding of *Salmonella* in chickens. Effective protection can be conferred during the first few days of a chick’s life regardless of time point and traditional hatchery delivery systems are sufficient to alter the microbiome and transfer donor material. Early microbiota intervention in chickens is a promising route of pathogen control in broiler flocks in the fight to control food-borne outbreaks in the human population.

## Introduction

Gastrointestinal intestinal infections caused by pathogenic bacteria such as *Salmonella* are one of the leading causes of foodborne illness worldwide. Principally, these infections are associated with the consumption of products from the poultry industry and as such control at the farm level is increasingly important (Chambers and Gong, 2011). Previously, farmers sought to increase feed efficiency and control pathogen infection by using antibiotic feed additives (Bedford, 2000, Castanon, 2007). However, the global trend in banning these additives has opened a niche market to find and develop new treatments that will prevent pathogen invasion and colonisation (Bedford, 2000, Chambers and Gong, 2011). These treatments should aim to prevent increases in antibiotic-resistant bacteria and help to balance the gastrointestinal (GIT) microbiota. One such method to achieve this is the use of GIT microbiota transplantation.

Manipulation of the GIT microbiota has been increasingly used in humans and animals to help combat varying diseases and infections (Khoruts and Sadowsky, 2016, Zhang *et al*., 2018). The acquisition of a complete and balanced microbiome early in life has been shown to help develop and strengthen the immune system, provide essential nutrients to the host and lead to overall better health in humans and animals. Historically, Faecal microbiota transplants (FMT) have been recorded as far back as the fourth century in China where they were used to treat patients with diarrhoea (Valiquette and Laupland, 2013). In humans, they have since been used to treat a range of infectious and communicable diseases thereby extending the range of applications (Gupta *et al*., 2016). In the food production industry, FMTs have been used, most often, to treat diarrhoea in animals from cows to pigs (Diao *et al*., 2016, Brunse *et al*., 2019, Diao *et al*., 2018, Kim *et al*., 2021, Islam *et al*., 2022). The broiler industry is distinct from other food production systems as chicks are generally hatched in large sterile industrial hatching incubators before being transferred to broiler grower units. This represents a unique environment in which chicks generally never encounter adult birds and thereby develop a rudimentary microbiota initially consisting of largely environmental microbes (Richards *et al*., 2019, Richards-Rios *et al*., 2020a).

Recently, Gilroy *et al*. (2018) showed that early manipulation of the microbiota via cecal microbiota transplant (CMT) in Ross 308 broiler chicks leads to lower intestinal colonisation of birds with the pathogen *Campylobacter jejuni*. A seeder bird infection model showed that *C. jejuni* transmission across groups was reduced when compared to non-transplanted infected birds (Gilroy *et al*., 2018). Alternatively, Richards-Rios *et al*. (2020b) demonstrated that the application of adult cecal content to the surface of eggs was sufficient in transferring elements of the microbiota to chicks which resulted in acceleration of microbiota development in chicks (Richards-Rios *et al*., 2020b). In parallel, Ramírez *et al*. (2020) established that cecal microbiota transplant and environmental transplants that are administered at day of hatch lead to a reduction in pathogen colonisation by either *Salmonella* spp or *C. jejuni*. The authors also investigated whether successional changes in the microbiota would lead to the establishment of a stable microbiota and found that one transplant from 14 days old birds into newly hatched chicks was enough to stabilise the GIT microbiota of broiler birds (Ramírez *et al*., 2020). Similarly, Zenner *et al*. (2021) showed transfer of a maternal microbiota to newly hatched chicks via passive colonisation resulted in increased gut microbiota diversity accompanied by increased levels of IgA and IgY when compared to birds kept under strict hygienic conditions (Zenner *et al*., 2021). Li *et al*. (2022), used an alternative approach to CMT by assessing the effect of Hen raising on the establishment of the microbiota. The authors showed that microbiota was transferred between hens and chicks which enabled the establishment of a balanced and diverse microbiota subsequently improving the stability following viral infection with H6N2 (Li *et al*., 2022). All these studies provide evidence that early life manipulation of the GIT microbiota does indeed lead to beneficial phenotypes in growing and adult birds.

Typically, when hatcheries send “Day-old” chicks to grower farms the birds will be between 4 and 72hrs of age, therefore any treatment or intervention utilising microbiota transplants should be effective across this age range. Hatcheries also deal with large numbers of birds hatching daily and therefore delivery methods for therapies should cover larger numbers of birds easily. The following study aims to address these two aspects presenting results from two trials used to assess the timing and route effects on microbiome acquisition and subsequent pathogen colonisation. The effect of transplanting CMT into newly hatched chicks between 4 and 72 hours of age and how this impacts colonisation and intestinal invasion with the *Salmonella enterica* serovar Typhimurium (ST4/74) is discussed. Early microbiota transplant in these birds prevents colonisation and invasion with ST4/74 and the timing of the transplant has a minimal effect on this inhibition. Using 16S rRNA analysis revealed distinct differences between the microbiota of CMT treated birds compared to PBS control birds. These findings present additional evidence that early life microbiota intervention can convey protective effects against *Salmonella* infection dynamics in broiler birds.

## Material and Methods

### Bacterial strains and culture conditions

*Salmonella enterica* Typhimurium 4/74 (ST4/74) was cultured from frozen stocks onto Luria Bertani (LB) agar for 24 hrs at 37°C with liquid cultures grown for 24 hrs in 10ml LB broth at 37 °C and then adjusted in fresh LB to a desired concentration of 10^6^ colony forming units per ml (cfu ml^-1^). All microbiological media were purchased from Lab M Ltd. (Heywood, Lancashire, United Kingdom).

### Cecal microbiota preparation

Cecal microbiota transplant (CMT) material was obtained from three pathogen free 8-week-old Ross 308 birds that had been reared in bio-secure conditions. Birds were euthanised via cervical dislocation prior to aseptic removal of the blind ended ceca. Cecal content were collected in sterile 50 ml falcon tubes and snap frozen in liquid nitrogen until processing. Cecal contents were then diluted 1:20 in sterile Phosphate Buffered Saline (PBS) supplemented with 10% Glycerol (PBS/glycerol-10) and filtered using a 70 μM cell strainer and stored in 2 ml aliquots at −80 °C until use.

### Experimental animals

All work was conducted in accordance with United Kingdom (UK) legislation governing experimental animals under project license PPL 40/3652 and was approved by the University of Liverpool ethical review process prior to the award of the licenses. All animals were checked a minimum of twice-daily to ensure their health and welfare was maintained. Two separate trials were performed to assess the effect of Timing (Trial A) and Delivery Route (Trial B) on the colonisation and faecal shedding of ST4/74 in Broiler chickens. Trial A: Embryonated Ross 308 eggs were obtained from a commercial hatchery and incubated in an automatic roll incubator under standard conditions for chicken eggs. Chicks were removed from the incubator post-hatch, split into four different groups and administered a 0.1 ml inoculum of CMT via oral gavage within 4, 24, 48 or 72 hours of hatching. Two additional control groups were given a 0.1 ml inoculum of PBS/glycerol-10. Trial B: Embryonated Ross 308 eggs were obtained from the same commercial hatchery as in Trial A and incubated as above. Chicks were removed from the incubator post-hatch split into 3 groups and either administered a 0.1 ml inoculum of CMT via oral gavage within 4 hours or at 24 hours post hatch sprayed with either CMT alone or CMT mixed with CevaGel droplet technology used for vaccine delivery. Control groups were given a 0.1 ml inoculum of PBS/glycerol-10. Chicks were housed in the University of Liverpool high-biosecurity poultry unit. Briefly, chicks were evenly distributed across different pens in climate-controlled rooms. Each pen used a wood shaving substrate for bedding and food and water were provided *ad libitum*. Up to 14-days post hatch (d.p.h) chicks were fed a pelleted vegetable protein-based starter diet and then from 14-d.p.h a pelleted vegetable protein-based grower diet was provided until the end of the experiment (Special Diet Services, Witham, Essex, UK). The Nutritional composition of the starter and grower diets is displayed in Table 1. Due to the high biosecurity levels maintained in the unit no coccidiostats or antimicrobials were added to either of the diets provided.

**Table 1.** Composition of Starter and Grower diets fed to birds *ad libitum* throughout each trial

| Analytical constituents (%) | Diet |  |
| --- | --- | --- |
|  | Starter | Grower |
| Crude fat | 2.7 | 2.4 |
| Crude protein | 18.9 | 15.6 |
| Crude fiber | 3.8 | 4.1 |
| Crude ash | 6.6 | 5.6 |
| Lysine | 0.99 | 0.69 |
| Methionine | 0.44 | 0.27 |
| Calcium | 1.05 | 0.89 |
| Phosphorus | 0.7 | 0.62 |
| Sodium | 0.15 | 0.15 |
| Magnesium | 0.17 | 0.22 |
| Copper | 15 mg/kg | 16 mg/kg |

### Effect of cecal microbiota transplantation on ST4/74 infection

During both Trial A and B, at 7-d.p.h a small number of birds from each group (minimum of n=10) were culled and cecal content were collected and snap frozen for 16S rRNA gene sequencing analysis. All birds in each group were weighed at 3-, 7-, 10-, 14-, 17-, and 21-days post hatch. At 7-d.p.h, all chicks in each experimental group were orally infected with 10^6^ CFU ml^-1^ of ST4/74 in 0.1 ml of Luria Bertani (LB) broth (Figure 1). At 3-, 7-, 10-, and 14-dpi cloacal swabs of all birds were taken to assess faecal shedding of ST4/74. At 3-, 7- and 14-dpi a small group of birds in each group (minimum of n =10) were culled via cervical dislocation for post-mortem sample collection.

**Figure 1:**
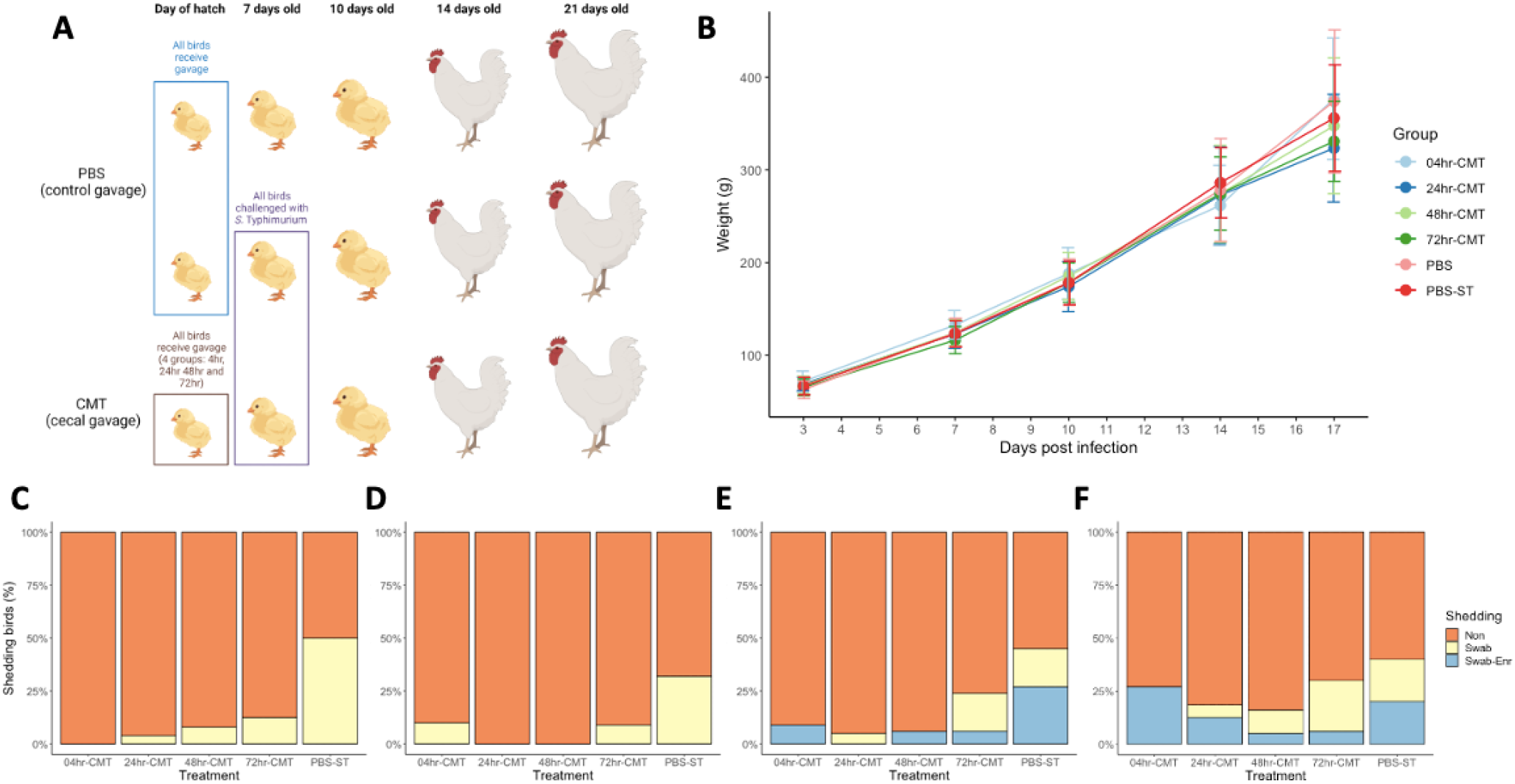
Timing effect study design, bird weights and ST4/74 shedding results. A) schematic of the transplant study showing birds given oral gavage of PBS at day of hatch and oral gavage of CMT within 4, 24, 48 and 72 hours of hatch. Time points show procedures carried out on groups; at 7 days old ST4/74 was given to one PBS group and all CMT groups with 10 birds from each group culled for post-mortem analysis and collection of cecal content for 16S rRNA analysis. At 10, 14 and 21 days old a subset of birds from each group were culled for post-mortem analysis and sample collection. All birds in each group were swabbed at 3, 7, 10 and 14 dpi to assess faecal shedding of ST4/74. Schematic produced using BioRender. B) All birds in each group were weighed at 3, 7, 10, 14, 17 and 21 days of age. There were no significant differences between groups on the average weight of birds. C-F) Results of faecal shedding of ST4/74 at 3, 7, 10, and 14 dpi as measured by cloacal swabbing and plating on to ST4/74 selective agar. Shedding was recorded as no shedding (Non, orange), ST4/74 detected following enrichment of swab (Swab-Enr, blue) and ST4/74 detected straight from swab onto plate (Swab, yellow). Compared to PBS treated groups, CMT treated groups were shown to have less faecal shedding of ST4/74 throughout the experimental period.

### Post-mortem analysis and sample collection

At post-mortem analysis, the spleen was located and placed in a sterile container and a portion of the liver was collected and placed in a separate sterile container, both to be used for downstream bacteriology. Both blind ended ceca were located and removed, and the contents manually expressed into a sterile container. The contents from one caecum were used for host bacterial enumeration and the contents from the other used for 16S rRNA gene sequencing.

### Assessment of *S. Typhimurium* load

To determine the level of ST4/74 intestinal colonisation within each group the collected cecal content was diluted in 9 volumes of PBS. Next, 10-fold serial dilutions were performed in PBS. Using the Miles & Misra method triplicate 20-ul spots were plated onto Brilliant Green Agar (BGA, ThermoFischer Scientific). Plates were then incubated aerobically for 24 h at 37 °C. *Salmonella* colonies were enumerated to give colony forming units per gram of cecal content (CFU/g).

### Assessment of *S*. Typhimurium shedding

At each time point cloacal swabs from all birds in each group were taken and plated onto BGA plates. The swab was then placed in universals containing 2 ml of *Salmonella* ONE broth (ThermoFischer Scientific). Plates and cultures were placed in aerobic incubators at 37 °C for 24 hours before being checked for growth. Enrichment in *Salmonella* ONE broth was plated onto fresh BGA plates and incubated for a further 24 hours. Shedding was recorded as scores based on whether *Salmonella* was detected and at which point of culture *Salmonella* was detectable (2 = direct from swab, 1 = following 24 h enrichment, 0 = not detected).

### DNA extraction and 16S rRNA analysis

DNA was extracted from each sample using the Zymobiomiccs DNA minikits (Cambridge Bioscience, UK) according to the manufacturer’s instructions. DNA was extracted from approximately 200 mg of cecal content. Initially, bead-beating was performed using a Qiagen TissueLyser at 30 Hz for 10 min. DNA was extracted from samples at each time point serially to ensure that storage time was equal. For each extraction, two controls were included; a blank extraction for contamination control and 75 μl of Zymobiomics Microbial Community Standard (Cambridge Bioscience, UK) to control for variations in DNA extraction efficacy. Extracted DNA was quantified using a Qubit dsDNA HS fluorometric kit (Invitrogen). Extracted DNA was then submitted to Centre for Genomic Research (University of Liverpool) for paired-end sequencing using the Illumina MiSeq platform. The V4 hypervariable region was amplified for 25 cycles to yield an amplicon of 254 bp using the primers in Table 1. Raw data files were trimmed for the presence of Illumina adapter sequences using Cutadapt version 1.2.1. Reads were further trimmed with Sickle version 1.200 with a minimum quality score of 20, any reads shorter than 15 bp after trimming were removed. QIIME2 (version 2020.11) was used for analysis of the Illumina data (Hall and Beiko, 2018). Amplicon sequence variants (ASVs) were assigned using the dada2 plugin (Callahan *et al*., 2016) and ASV tables were produced and exported in biological observation format (BIOM) for individual timepoints from each trial. Taxonomy was assigned using the q2-classifier plugin to generate a taxonomy table for each ASV. The classifier was trained using the 99% green genes dataset using the primer sets used for amplification of the V4 region during sequencing. The taxonomy table, ASV table and rooted phylogenetic tree, along with the metadata file were exported for use in further analysis.

### Statistical analysis and data summaries

Community analyses using the exported taxonomy and ASV tables were performed in RStudio version 4.1.3 using the Phyloseq (McMurdie and Holmes, 2012, McMurdie and Holmes, 2013) and Vegan packages. Briefly, Alpha and Beta diversity were performed at a sequencing depth of 7000. With Alpha diversity measured with an observed and Shannon metric. Beta diversity was measured using Bray-Curtis dissimilarity matrix and used to draw PCoA plots. Top 30 most abundant taxa at each time point were identified and Kruskal Wallis test was used to compare groups to determine differences.

## Results

### Early-Life cecal microbiota transplantation reduces faecal shedding of ST4/74 irrespective of delivery time

A simple oral gavage model of infection was used to assess ST4/74 colonisation and shedding in broiler birds that were orally gavaged with either PBS or CMT in the first few days of life (Figure 1A). Birds were weighed regularly throughout the experiment to determine if there were any effects of transplant on this measure of flock performance. Figure 1B shows that there were no significant differences between the average weight of birds between groups throughout the experiment. Monitoring of faecal shedding of *S*. Typhimurium 4/74 showed a markedly reduced effect across birds that received transplant compared to birds that received vehicle only (Figure 1C-F). At 3 dpi faecal detection of *S*. Typhimurium was greatest at 49% in the PBS challenged group (Figure 1C) with no detectable ST4/74 shed in the faeces of birds given CMT at 4hr post hatch. There is a small increase in the number of birds shedding *S*. Typhimurium in the faeces in birds given CMT at 24, 48 and 72hrs post hatch but this number is markedly reduced compared to PBS controls. At 7dpi, faecal shedding was detected in around 10% of birds in the 4hr and 72hr CMT groups, but not in the 24hr or 48hr CMT groups. By 7dpi there are fewer PBS control birds detected as shedding 4/74 compared to 3 dpi but this is still more than CMT groups (Figure 1D). By 10 and 14 dpi all groups had birds with detectable *S*. Typhimurium shed in the faeces however, the CMT groups still had fewer shedding birds compared to PBS controls

### Intestinal colonisation of ST4/74 is reduced in CMT transplanted birds compared to controls

At 3, 7 and 14 dpi. A subset of birds from each group were taken for post-mortem analysis to assess the bacterial load of ST4/74 in the ceca, liver and spleen. At 3, 7 and 14 dpi colonisation of the ceca with ST4/74 was significantly reduced across all treatment groups (Figure 2). At 3 dpi, 7 birds in the PBS control group showed high levels of colonisation with ST4/74 with only 2 birds in the 24hr group and 1 bird in the 48hr group showing colonisation of the ceca (Figure 2A). Figure 2B shows that at 7 dpi there were 5 birds in the PBS group with ST4/74 colonisation in the cecal content with the 24 and 48hr groups both having only 1 bird with detectable ST4/74. The 72hr group had 3 birds positive for ST4/74 in the ceca but the level was lower than PBS controls. Birds in the 4hr group remained uncolonized. By 14 dpi 4 birds were shown to be positive for ST4/74 in the PBS and 48hr groups all other groups remained negative for colonisation of the ceca (Figure 2C). Throughout the experiment ST4/74 was unable to be detected in the liver and the spleen of birds in all groups.

**Figure 2:**
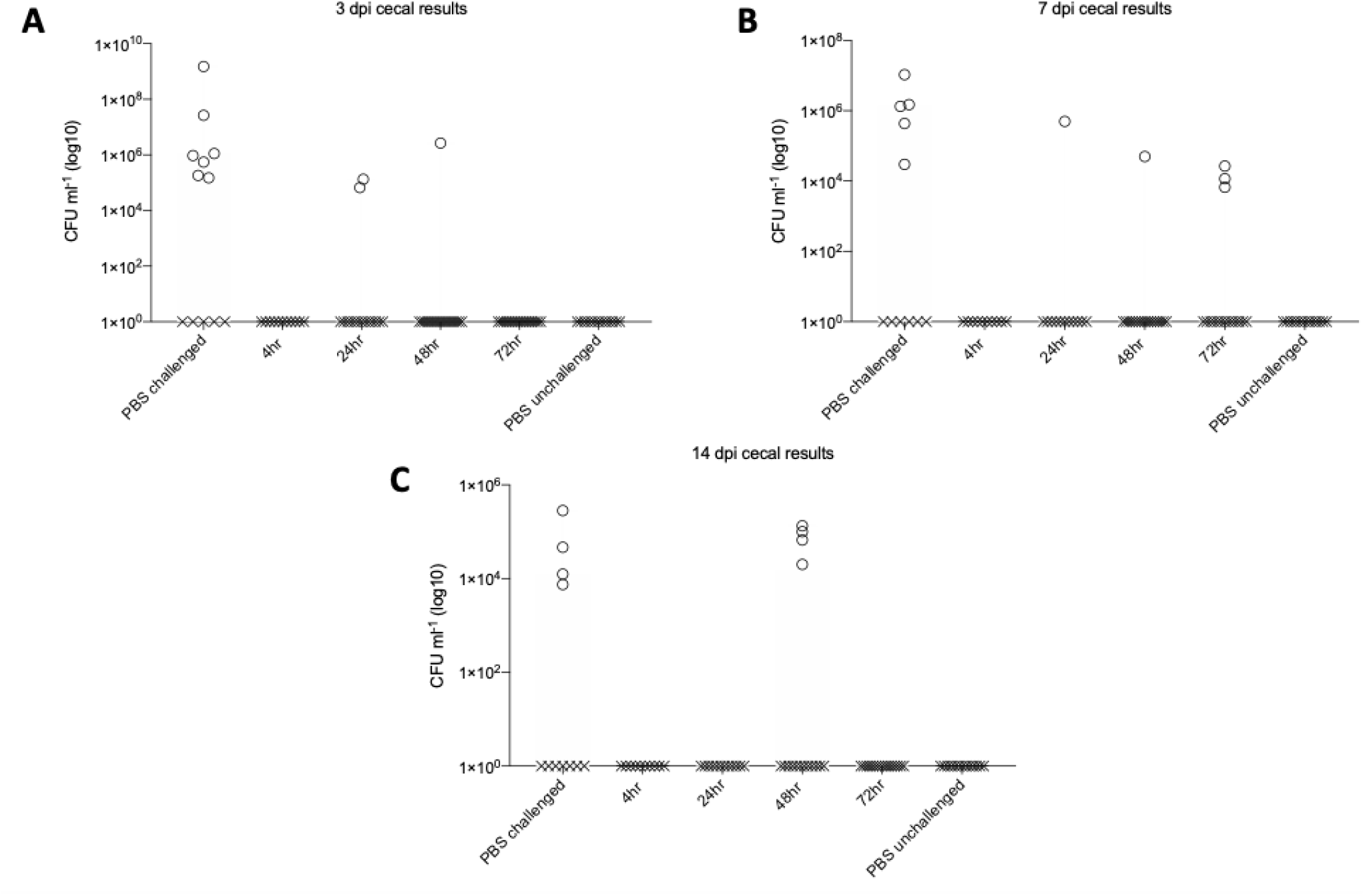
Colonization of cecal content of birds with ST4/74. A) At 3-dpi over 50% of the birds in the PBS challenged group showed detectable numbers of ST4/74 colonizing the cecal content. Of the CMT groups two birds in the 24hr group and 1 bird in the 48hr group showed detectable levels of ST4/74 in the cecal content. All other groups did not appear to be colonized by ST4/74. B) At 7-dpi three of the transplant groups (24, 48 and 72) had birds which showed colonization in the cecal content however with a lower CFU ml^-1^ compared to PBS challenged controls. C) At 14 dpi 4 birds in the PBS challenged and 48hr CMT group were colonized at similar levels with ST4/74.

### Effect of CMT delivery route on ST4/74 faecal shedding in broiler chicks

In a commercial setting it would be impractical to use oral gavage as a mode of delivery for any therapy. Therefore, delivery of CMT via spray or a commercial gel droplet product (CevaGel) was also investigated (Figure 3). As compared to the previous trial there were no significant differences in the bird weights between experimental groups (Figure 3B). We were able to show that CevaGel delivery appeared to offer the greatest level of protection against faecal shedding of ST4/74 in the birds with less than 10% of birds detected as shedding ST4/74 at each time point throughout the study (Figure3 C-E). Gel delivery seemed to be the least effective method with almost 70% of the birds shown to be shedding ST4/74 in the faeces by 14 dpi. Despite this ST4/74 was only detected in this group following enrichment potentially suggesting low numbers of CFU ml^-1^. Further analysis would be required to determine if this level of colonisation would be transmissible and infectious. Across most time points assessed, PBS and 4hr-CMT groups appeared to be comparable in detection of ST4/74 in the faeces. Although, most if not all the birds were only shown as ST4/74 positive in the 4hr-CMT group following enrichment of the swab, again suggesting a lower number of CFU ml^-1^ compared to the PBS challenged group.

**Figure 3:**
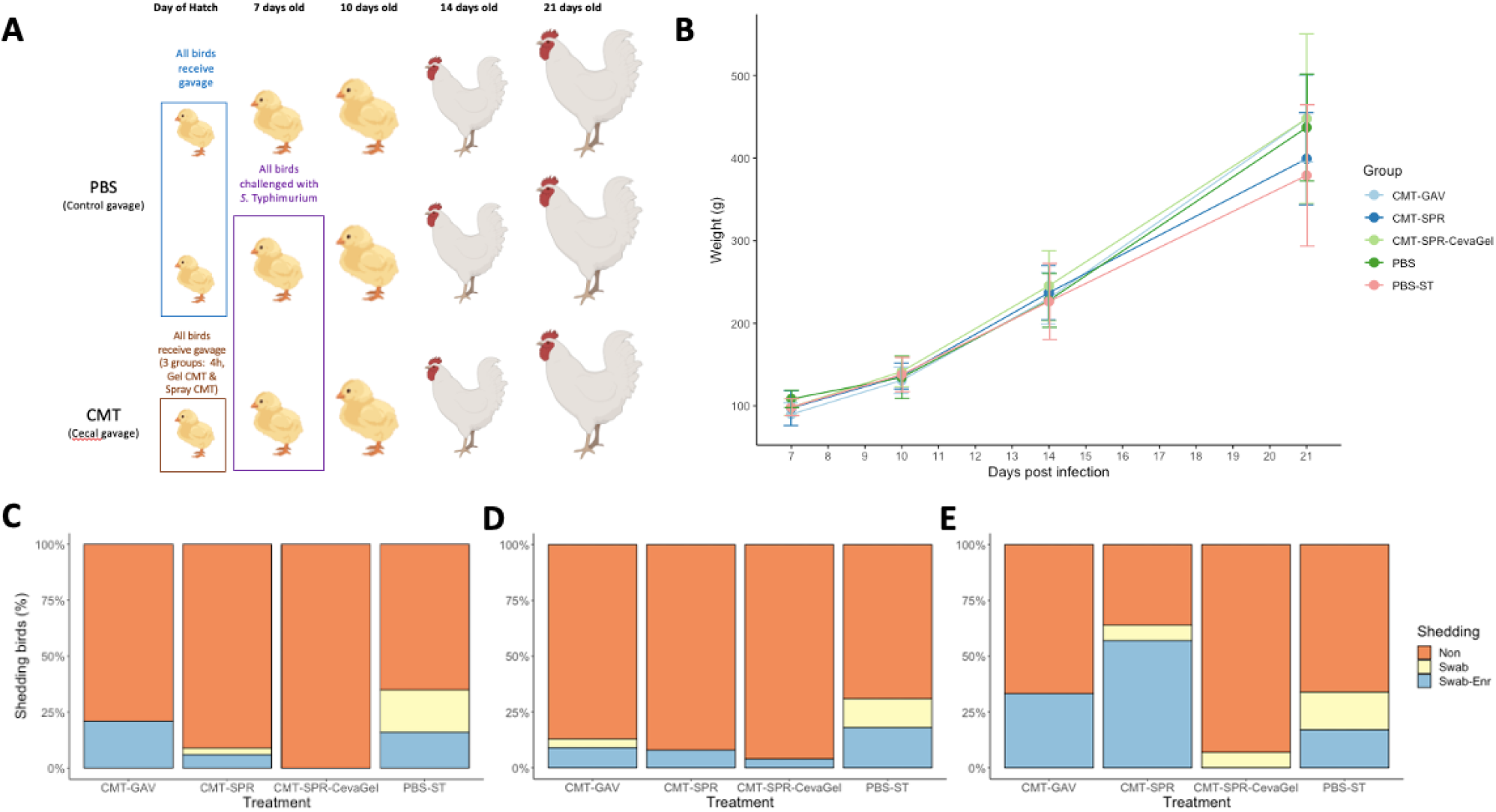
Route effect study design, bird weights and ST4/74 shedding results. A) schematic of the transplant study showing birds given oral gavage of PBS at day of hatch and oral gavage of CMT within 4 hours of hatch. Two groups of birds were given transplant in the form of spray delivery (CMT-SPR) or gel spray (CMT-SPR-CevaGel) delivery within 24 hours of hatch. Time points show procedures carried out on groups; at 7-days old ST4/74 was given to one PBS group and all CMT groups with ∼10 birds from each group culled for post-mortem analysis and collection of cecal content for 16S rRNA analysis. At 10-, 14-and 21-days old a subset of birds from each group were culled for post-mortem analysis and sample collection. All birds in each group were swabbed at 3-, 7-, 10- and 14-dpi to assess faecal shedding of ST4/74. Schematic produced using Biorender. B) All birds in each group were weighed at 3-, 7-, 14- and 21-days of age. There were no significant differences between groups on the average weight of birds. C) Results of faecal shedding of ST4/74 at 3,- 7-, and 14-dpi as measured by cloacal swabbing and plating on to *Salmonella* selective agar. Shedding was recorded as no shedding (Non, orange), ST4/74 detected following enrichment of swab (Swab-Enr, blue) and ST4/74 detected straight from swab onto plate (Swab, yellow). Compared to PBS treated groups, CMT treated groups were shown to have less faecal shedding of *Salmonella* throughout the experimental period.

### Microbiome analysis reveals distinct differences between CMT groups and PBS controls

#### CMT groups have greater species richness compared to PBS controls

In general, at 7-dph and 3-dpi birds in the CMT groups had greater numbers of observed species as well as higher species richness compared to PBS groups across both trials (Figure 4 & 5). By 7-dpi and 14-dpi the alpha diversity appears to even out between the CMT groups, and the PBS challenged groups. Interestingly, during the timing trial birds from the PBS challenged group seem to have a quicker development in microbiota compared to PBS unchallenged birds (Figure 4). At 3-dpi (Figure 4B), 7-dpi (Figure 4C) and 14-dpi (Figure 4D) the species richness in the PBS-challenged birds gradually becomes more comparable to the CMT groups compared to PBS-unchallenged groups. During the Route trial the microbiomes of the birds in the CMT and PBS groups are comparable for species richness by 3-dpi (Figure 5B) and remain so up to 14-dpi (Figure 5D).

**Figure 4.**
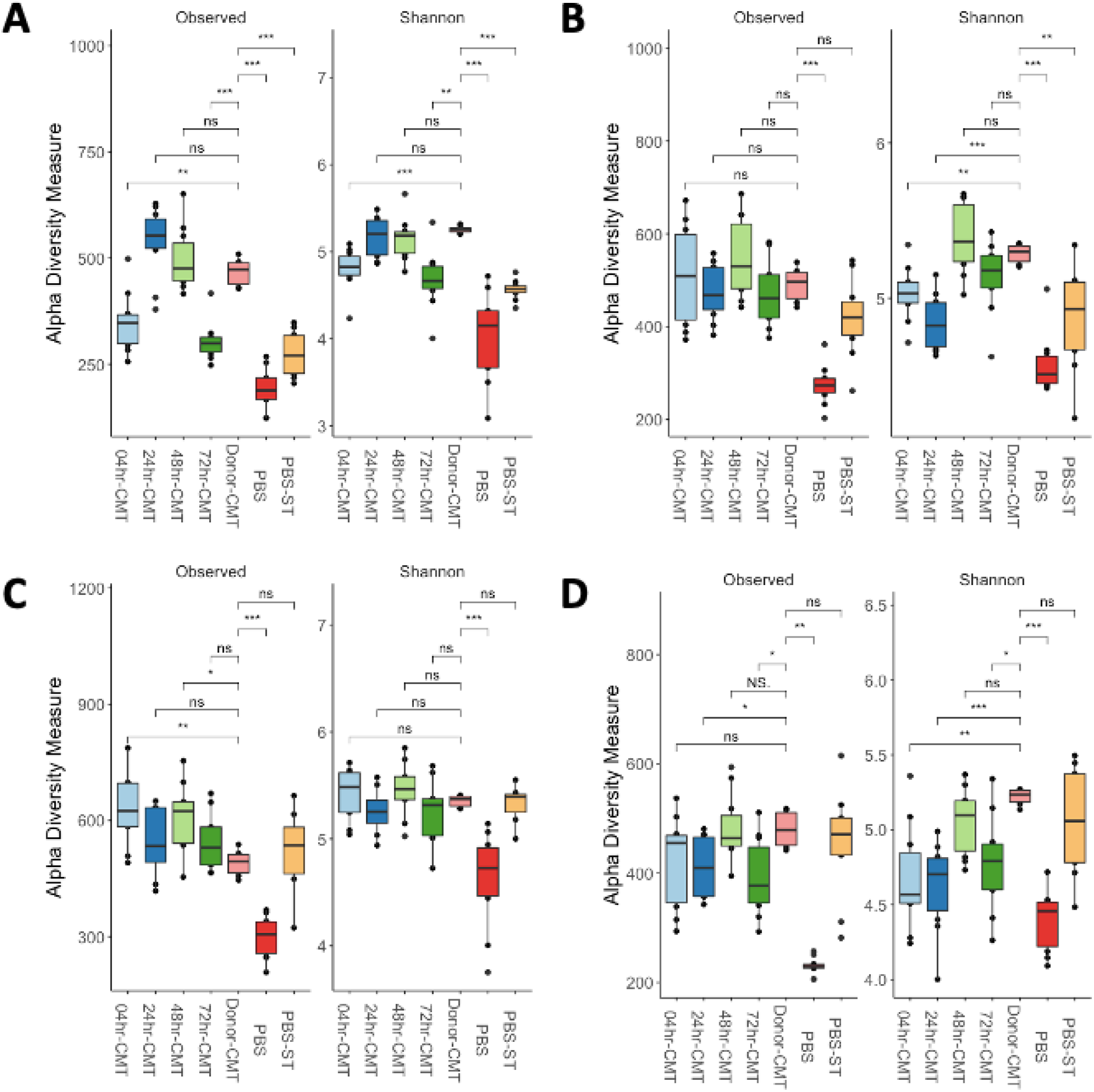
Alpha diversity measures comparing timing effects on microbiome richness. Alpha diversity was calculated using Observed and Shannon measures at a sequencing depth of 7000. A) At 7-dph and prior to infection with ST4/74 there were some differences noted in the observed species compared with donor-CMT. The CMT groups showed higher alpha diversity like the donor-CMT when compared with PBS groups. B) At 3-dpi there is no significant difference in observed species of the CMT groups compared to the donor-CMT. Shannon diversity shows that CMT groups are still higher than PBS groups however alpha diversity of PBS-ST group appeared to be developing quicker than just PBS. C) By 7-dpi the PBS-ST group was now as diverse in species abundance as the CMT and donor-CMT groups compared to PBS alone. D) AT 14-dpi there is still higher diversity in the PBS-ST and the CMT groups compared to PBS alone groups suggesting that PBS group microbiota has not developed as quickly as the infected group.

**Figure 5:**
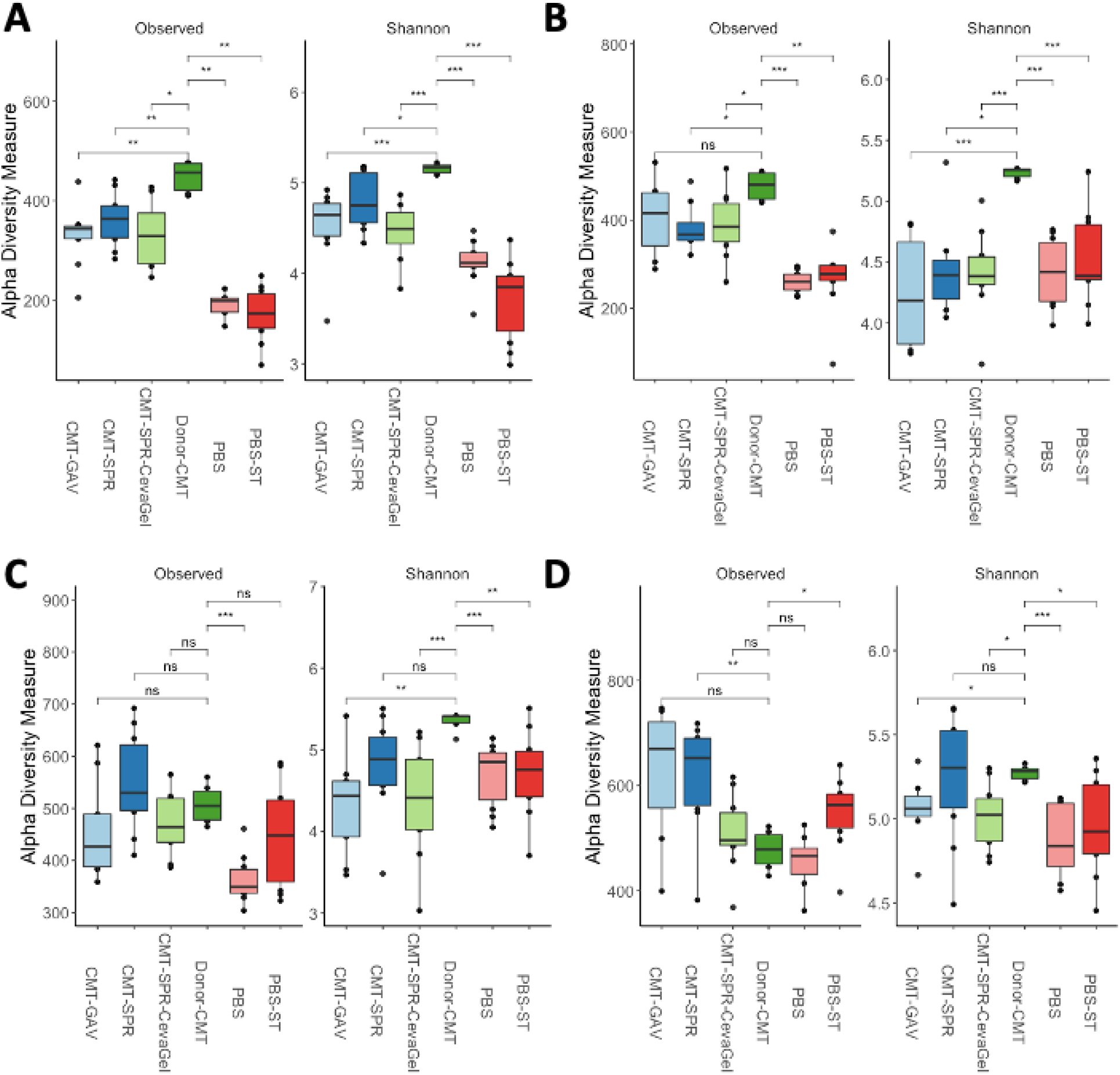
Alpha diversity measures comparing route effects on microbiome richness. Alpha diversity was measured using Observed and Shannon measures at a sequencing depth of 7000. A) At 7dph the CMT transplant groups have comparable numbers of observed species and richness and are closer to the donor material than the PBS groups, but the donor material still has more species and greater richness than all the trial groups. B-D) From 3-14-dpi the alpha diversity of the CMT groups compared to the PBS groups does not have as distinct differences as in the timing trial. Species richness between the groups is comparable and up to 14-dpi is significantly lower than that of the donor material.

#### CMT treatment significantly affected microbial composition

When measured with Bray-Curtis dissimilarity index clear separation of CMT groups to PBS groups could be detected. Figure 6 A shows that there is distinct separation between the PBS control and CMT groups at 7-dph. The CMT groups cluster much closer to each other and the donor CMT material than they do with the PBS groups showing that early-life intervention of the microbiome has altered the initial microbiome in chicks. Interestingly, at each of the timepoints assessed post-infetion with ST4/74 the PBS-challenged group begins to cluster more closely with the CMT groups (Figure 6B-D). The PBS-unchallenged group remained separated at every time point (Figure 6B-D). During the route trial the clustering of the microbiomes does not appear to be as strong as during the timing trial however a similar pattern of clustering was observed (Figure7). At 7-dph, the PBS groups cluster close together with the CMT groups clustering together and more closely to the donor material than the PBS groups (Figure 7A). Again, following infection with ST4/74, the micobiomes of the PBS-challenged groups start to cluster more closely with the CMT groups as compared to the PBS-unchallenged groups (Figure 7B-D).

**Figure 6.**
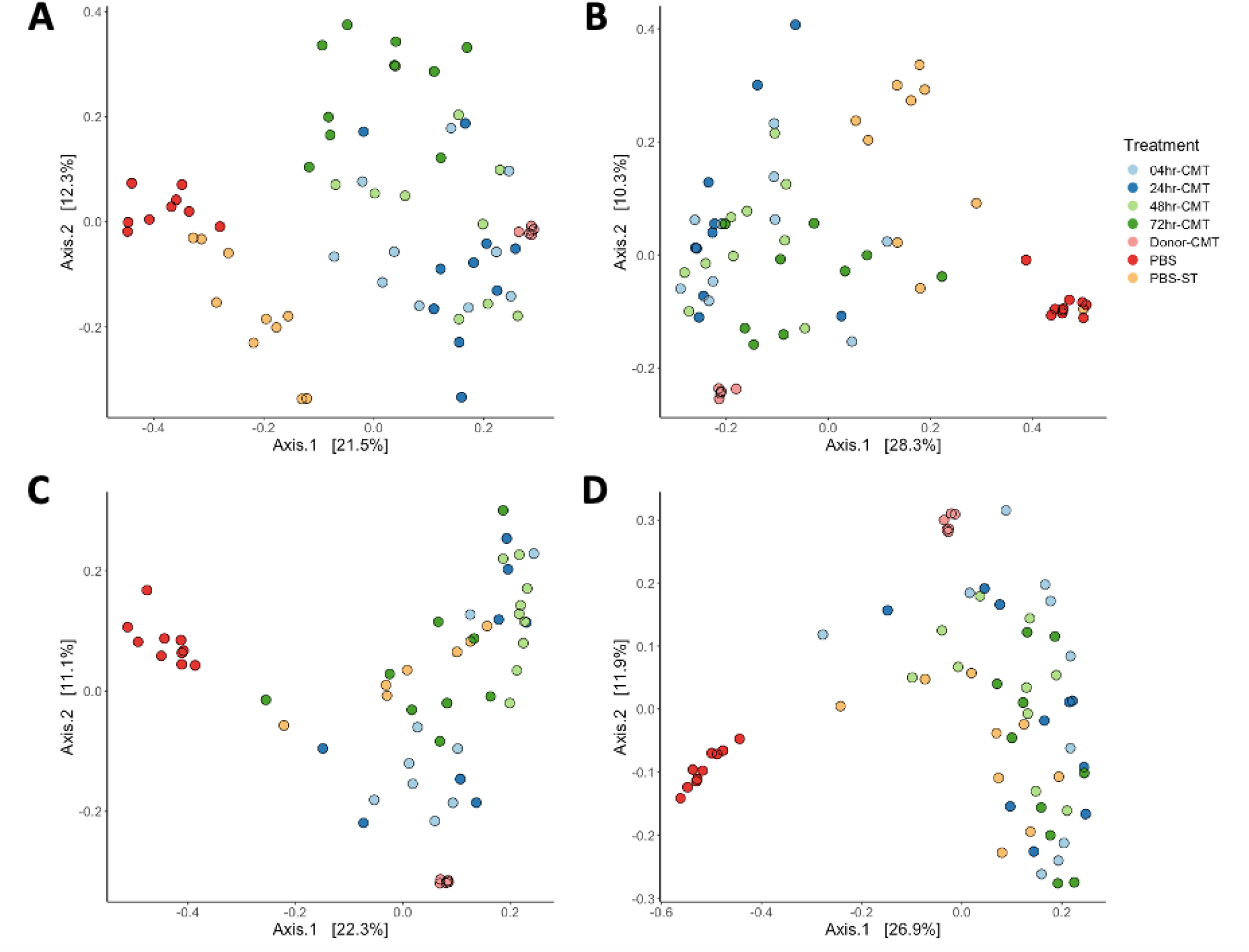
Effect of timing on patterns of community composition in CMT birds compared to PBS birds. Principle coordinate of Bray-Curits dissimilarity plots show distinct differences between the composition of CMT groups compared to PBS groups. At 7-dph (A) and prior to infection with ST4/74 the CMT groups cluster closely with the donor material whilst the PBS groups cluster closer together. From 3-dpi (B) shifts in the composition of the PBS-ST group but not the PBS group were observed. At 7-dpi (C) the PBS-ST groups had fully shifted to compositions similar to CMT groups and by 14-dpi (D) separation of the PBS group form the CMT and PBS-ST groups was most distinct.

**Figure 7:**
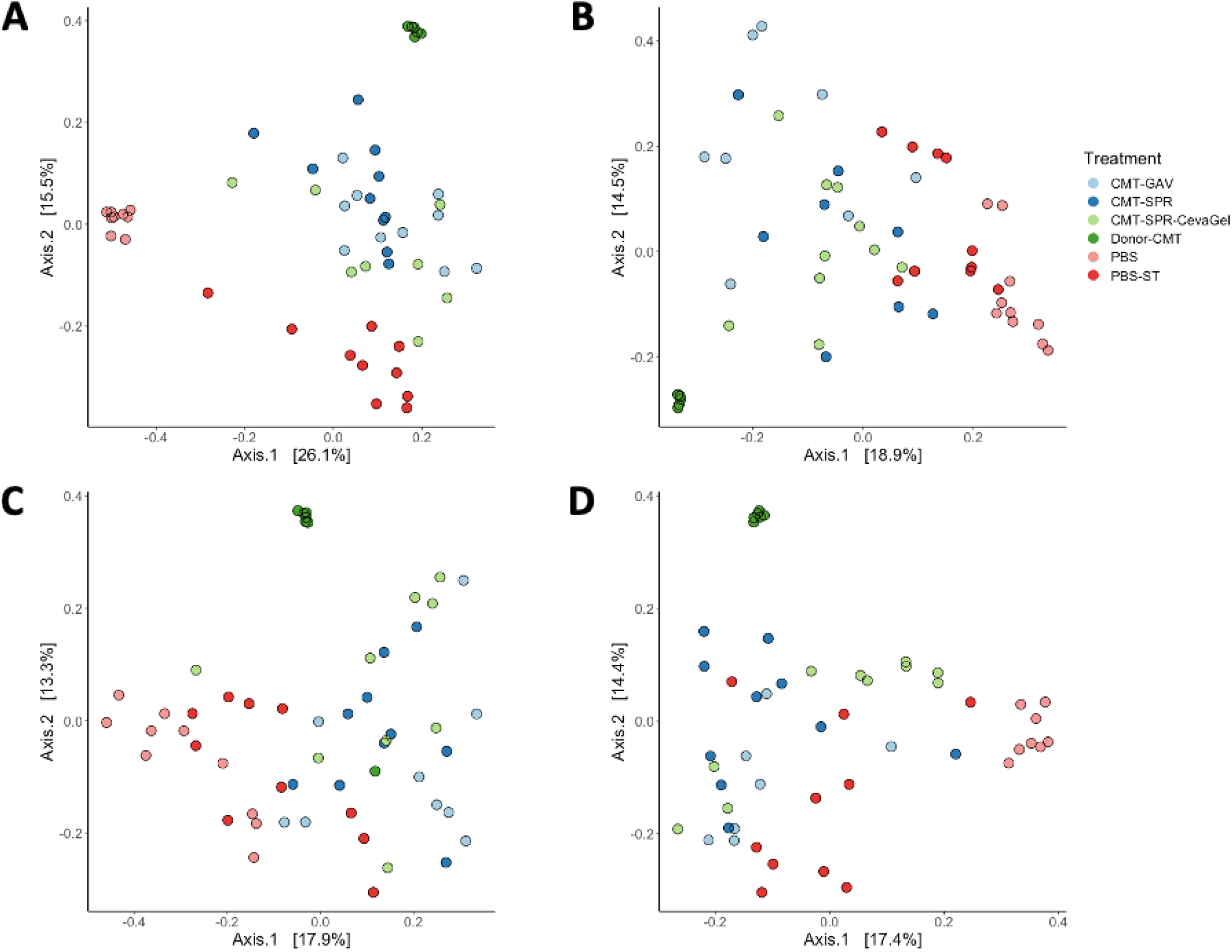
Effect of route on patterns of community composition in CMT birds compared to PBS birds. Principle coordinate of Bray-Curits dissimilarity plots show differences between the composition of CMT groups compared to PBS groups, though not as distinct as in the timing trial. At 7-dph the distinction between the CMT and PBS groups was most clear with the CMT groups appearing to cluster more closely with the donor CMT than the PBS groups. Similar shifts appear to occur with the PBS-ST group at 3-dpi (B) as occurred in the timing trial with the PBS-ST group developing a composition like that of the CMT groups compared to PBS groups. By 7-dpi (C) and 14-dpi (D) more of the PBS-ST groups cluster with the CMT groups than the PBS groups.

### The anti-inflammatory bacterium *Faecalibacterium prausnitzii* is significantly more abundant in CMT groups compared to controls

Finally, 16S rRNA analysis was used to determine if specific taxa from the microbiota could be attributed to the reduction in faecal shedding of ST4/74 in the CMT groups. The top 30 most abundant taxa from each time point were analysed to determine which bacteria, if any, may be contributing to these effects. *Faecalibacterium prausnitzii* was the only species in the Top 30 that was significantly more abundant in the CMT groups compared to PBS controls (Figure 8). During both the timing and the route trial the relative abundance of *F. prausnitzii* in the CMT groups compared to PBS controls is significantly higher at 7-dph (pre-infection, Figure 8A and 8E). Following infection with ST4/74 the relative abundance of *F. prausnitzii* starts to increase in the PBS challenged groups but not PBS unchallenged groups in both trials (Figure 8B-D, F and H). During the route trial at 7-dpi there is no significant difference in the relative abundance of *F. prausnitzii* in the CMT-SPR group and PBS-ST groups compared to the PBS unchallenged group (Figure 8G). *F. prausnitzii* is still significantly more abundant at this time-point in the orally gavaged and Gel spray CMT groups (Figure 8G).

**Figure 8:**
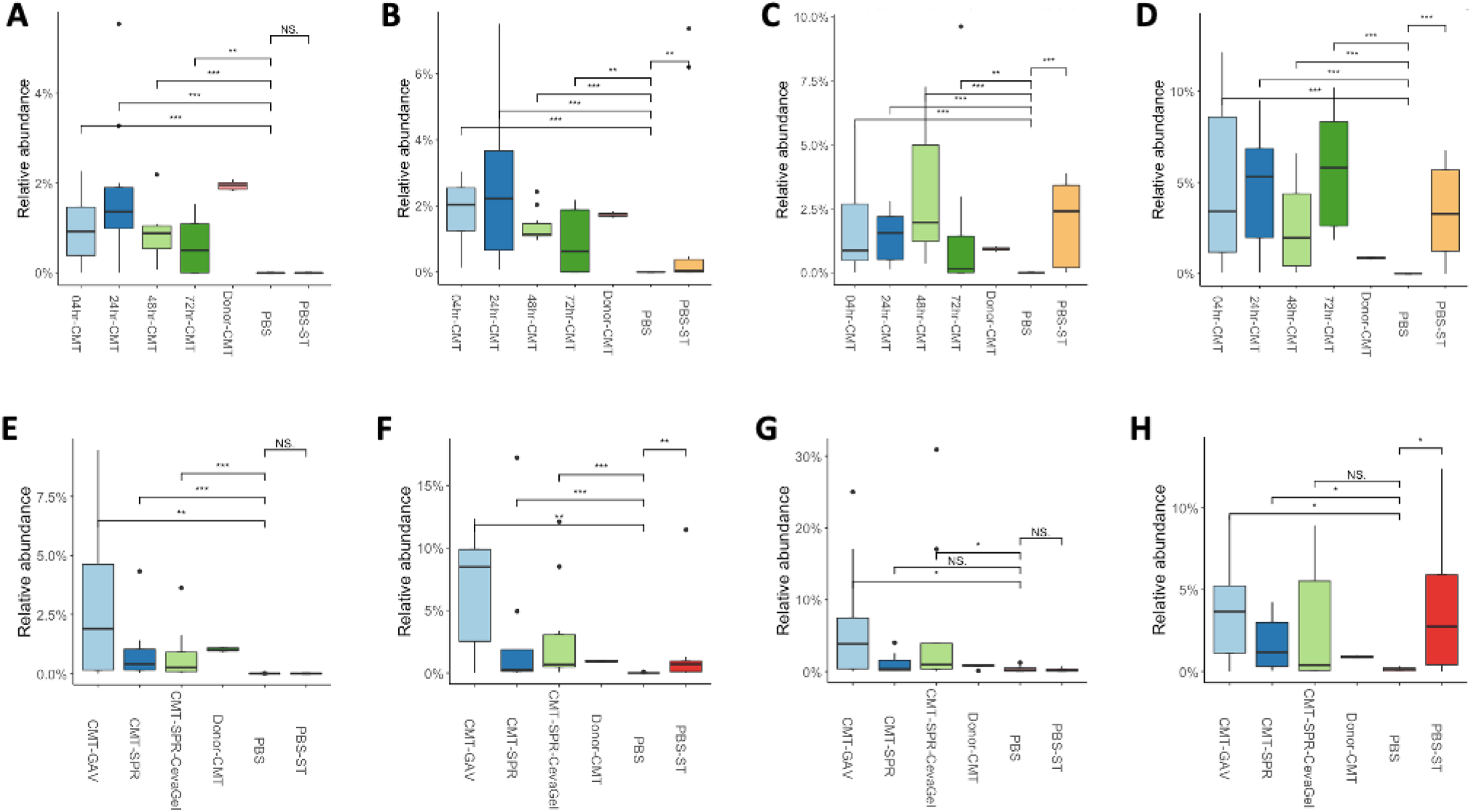
*Faecalibacterium prausnitzii* abundance is higher in CMT chicks than PBS controls both pre-infection and during the early stage of infections. The Relative abundance of *F. prausnitzii* in the chicken gut microbiome of broiler chickens in CMT groups vs PBS controls from Trial A (A-D) and Trial B (E-H). At 7-dph and prior to ST4/74 infection the relative abundance of *F*.*prausnitzii* is significantly higher in the CMT groups compared to the PBS consistent for both trials (A and E). Following infection, the relative abundance of *F. prausnitzii* increases in PBS groups that were challenged with ST4/74 but not in PBS unchallenged groups (B-D F and H). This is not apparent at 7-dpi during the route trial where there is no significant difference in the relative abundance of *F. prausnitzii* between the PBS groups or between the spray CMT group (G).

## Discussion

The present study aimed to influence the development and acquisition of the chicken gut microbiome early in life. Chickens in the commercial setting offer a unique opportunity to better assess the effects of microbiome manipulation compared to other food industry systems such as cattle, pigs, and sheep. Generally, the poultry meat or broiler industry is split into three sections: breeder flocks composed of birds used to produce eggs, hatcheries; where eggs are hatched in batches prior to transfer of chicks to the final section; grower farms, where they are reared until slaughter. In these production systems, a single flock will be approximately the same age and never be in contact with adult birds during the stages of microbiome acquisition. This can lead to delayed development of the microbiota. In fact, studies show that the initial microbiome of chickens from hatcheries is mostly composed of environmentally acquired bacteria (Pedroso *et al*., 2005, Stanley *et al*., 2014, Ballou *et al*., 2016, Donaldson *et al*., 2017, Richards-Rios *et al*., 2020b).

Previously, it has been shown that the early composition of cecal microbiomes in chicks show poor diversity and are comprised of mostly environmental species (Richards *et al*., 2019). It was demonstrated that by 21-dph the microbiota had matured and become stable across three different chicken breeds leading to the rationale within the current study to complete trials by 21-dph (Richards *et al*., 2019). The current study showed that transplantation of adult cecal content to chicks within the first few days of life leads to distinct differences in the composition of the microbiota when compared to PBS control birds. Birds from the CMT groups showed significantly more species and greater richness scores in the first week of life compared to PBS birds. These findings are consistent with that of others which also showed that administration of microbiota to chicks early in-life confer greater species diversity and differences in composition compared to control birds (Ramírez *et al*., 2020, Zenner *et al*., 2021, Glendinning *et al*., 2022). The present study showed that early-life intervention with a more complete microbiome confers a protective effect on birds subsequently challenged with *Salmonella* Typhimurium. In general, birds given CMT within the first 3 days post-hatch were shown to shed less *Salmonella* in the faeces compared to PBS control birds. Furthermore, if birds in CMT groups did have detectable *Salmonella* in the faeces it was more often detected following overnight enrichment of cloacal swabs. This potentially indicates that the *Salmonella* shed was of a lower CFU ml^-1^. Further work is required to determine if this low level of *Salmonella* would prove to be less transmissible and infectious within a flock compared to *Salmonella* levels which can be detected directly from a cloacal swab.

As described above, chickens are kept as single age cohorts throughout the production system. Generally, a batch of “day-old” chicks can be anywhere from 4hrs to 72hrs old, therefore the effectiveness of microbiome transplants needed to be assessed across these age ranges. Should microbiome transplants or more complex combinations or consortia of probiotics be utilised as an effective treatment option in the combat of *Salmonella* colonisation in the broiler industry they need to be robust and effective across the age ranges of the birds in each batch. We demonstrated that the timing of CMT administration had limited effect on the acquisition of the microbiome in treated birds and that *Salmonella* colonisation was reduced in all transplant groups compared to controls. The findings from this study are complementary to those of Varmuzova *et al*. (2016), who showed that prevention of *Salmonella* colonisation of birds is ineffective when using transplants from birds 1 week of age. However, when using birds between 3-42 weeks of age *Salmonella* colonisation is significantly reduced. Combined with the data shown in this study, development of the microbiota and manipulation during early stages has profound effects on subsequent pathogen invasion. Provision of more complete microbiotas during the first week of life impart clear benefits in older birds when challenged with a pathogen. However, Varmuzova *et al*. (2016) did show that this protection is only effective as a prophylactic treatment and not as a therapeutic, this was not tested in the current study.

Whilst oral gavage is an effective way administer transplants in an experimental setting this would not be possible/feasible in a hatchery setting. Therefore, to align with hatchery practices, traditional spray delivery methods such as those used to administer current vaccinations to chicks was tested to see if these were also effective at colonising and conveying protection to chicks. During a second trial the effect of spray delivery and Gel droplet technology on microbiota acquisition and *Salmonella* protection was compared to that of oral gavage. Whilst the results from this trial were not as strong as the results from the timing trial, distinct differences in the microbiota between CMT and PBS birds were observed. Of the two spray style delivery systems Gel appeared more effective at protecting birds from *Salmonella* colonisation with very low levels of *Salmonella* detected in faeces. Gel drop is a promising route of administration as the droplet technology prevents chicks from becoming ‘wet’ during vaccinations. This is key as it prevents chilling and stress as a result of vaccination (https://www.immucox.com/CevaGel-Droplet-Technology). Stress responses have also been shown to have adverse effects on the microbiota of chickens (U. Bello *et al*., 2018) and so limiting stressors during early life alongside administering a treatment to improve microbiota development may prove beneficial.

Interestingly, microbiome analysis revealed that infection with ST4/74 appeared to shift the microbiome composition of infected control birds as demonstrated by the rapid shifts in microbiome composition of PBS-ST birds compared to PBS birds (Figures 6, & 7). At 3-, 7-, and 14-dpi there were shifts in the Alpha and Beta diversity of PBS-challenged birds resulting in microbiome compositions like that of the CMT birds. These shifts were accompanied by a gradual reduction in colonisation of PBS-ST birds as well as reduced detection of *Salmonella* on cloacal swabs. As an enteric pathogen *Salmonella* species have developed numerous strategies to compete with resident microbiota, bypass colonisation resistance and cause infections in various hosts (Gart *et al*., 2016, Ahmer and Gunn, 2011). Induction of an inflammatory response in the gastrointestinal tract appears to be a key factor of benefit to *Salmonella* at the expense of the host microbiota (Chirullo *et al*., 2015, Drumo *et al*., 2015, Mon *et al*., 2015). However, it has been shown that higher community diversity of the microbiota in animals leads to less susceptibility to pathogens and pathobionts (Kamada *et al*., 2013). Specifically, inoculation of chicks with microbiota from chickens between 3 and 42 weeks of age significantly reduces colonisation of the ceca and liver with *S. enteritidis* (Varmuzova *et al*., 2016). Additionally, Pedroso *et al*. (2021) showed that *Salmonella* abundance decreased in experimentally infected birds as the species diversity of the microbiota increased. Birds were not given microbiota transplants for this study. Rather, they were infected with *Salmonella* and subsequently split into exclusive (*Salmonella* qPCR negative) or permissive (*Salmonella* qPCR positive) groups. It was also noted that specific species from the *Clostridiales* family were negatively associated with *Salmonella* abundance (Pedroso *et al*., 2021). Members of this family are known producers of Short Chain Fatty acids (SCFAs) such as butyrate, which is known to suppress type III secretion systems associated with *Salmonella* cell invasion. This could indicate a method of protection by microbiome transplantation (Van Immerseel *et al*., 2003, Gantois *et al*., 2006). Taken together the introduction of a more diverse community of microbes early in chick development could prove effective in preventing *Salmonella* colonisation in chickens thereby reducing the risk of *Salmonella* contamination in poultry meat.

Despite the positive effects noted from complete microbiome transplants in numerous studies, it may not be feasible to mass produce these highly complex communities. Therefore, it would be beneficial to identify specific species whose abundance or colonisation of the chicken GIT could confer protection against *Salmonella* infection alone or as part of a mixed probiotic. Pedroso *et al*. (2021), showed that the anti-inflammatory bacterium *Faecalibacterium prausnitzii* correlated with reduced *Salmonella* load. In the present study, significant differences in the abundance of *F. prausnitzii* between CMT and PBS groups were noted, suggesting this could be a key species involved in protection of chicks from colonisation with *Salmonella. F. prausnitzii* was higher in CMT birds compared to PBS birds and that its abundance increased in PBS-ST birds compared to PBS birds from 3-dpi. *F. prausnitzii* is known to provide benefits to the host by conveying anti-inflammatory effects in the GIT (Miquel *et al*., 2013, Wu and Wu, 2012). These anti-inflammatory effects have been shown to be mediated through the production of SCFAs such as butyrate (Gantois *et al*., 2006, Lenoir *et al*., 2020). Indeed, it has been shown that depletion of butyrate-producing bacteria from the GIT of chickens allows for the expansion of *Salmonella (Rivera-Chavez et al*., *2016)*. Butyrate supplementation of chicken feed has not only been proven to reduce the colonisation and faecal shedding of *Salmonella* (Wu *et al*., 2016), but also improve intestine development and provide growth advantages to poultry (Van Immerseel *et al*., 2005). In line with these studies, *in vitro* studies using chicken epithelial cells showed protection against invasion with ST4/74 using different concentrations of SCFAs (Butyrate, Acetate and Propionate) and filtered CMT (Supplementary figure 1). CMT performed as well as individual SCFAs however, data to show the SCFA composition of CMT was not collected. This should be performed in future studies to determine SCFA composition of CMT. It would also be beneficial to determine the direct interactions, if any between, *Salmonella* spp. and *F. prausnitzii* to further understand how these bacteria may be interacting within the host. Indeed, testing of a symbiotic-style product could be utilised, whereby *F. prausnitzii* is used as a probiotic supplement alongside butyrate-supplemented feed. Overall, *F prausnitzii* is increasingly being identified as a potential driver of protection in the chicken GIT and as a candidate for future probiotic supplementation in the fight to reduce *Salmonella* colonisation and infections in poultry.

In summary, it was shown that transplantation of the cecal microbiota into broiler chicks offers protection against *Salmonella* colonisation and subsequent faecal shedding. The timing of transplantation has minimal effect on the uptake and composition of microbiome compared to controls. Delivery route had minimal effect on microbiota acquisition however, spray delivery was least effective in reducing shedding of *Salmonella* compared to Gel-droplets and oral gavage. The effects of specific species and their effect on the colonisation and faecal shedding of *Salmonella* still needs to be determined however, higher abundance of *F. prausnitzii* in chicks pre-infection and the increase seen following infection in control birds could suggest a role for this bacterium in protection of chicks from intestinal colonisation. Future studies should assess the effect of *F. prausnitzii* supplementation in broiler chicks on *Salmonella* infections. This could be done either alone or in conjunction with other probiotic species or feed supplementation to potentially develop new probiotic/ synbiotic cocktails to protect chickens in the broiler setting. Fundamentally, this can lead to a reduction of *Salmonella* contamination in poultry meat with subsequent benefits of decreasing *Salmonella* induced gastroenteritis in the human population.

## Author Contributions

SP, AD and PW were supported by a BBSRC research grant (BB/R008914/1) and designed the study. SP, ALW, AW and SJ carried out experiments and monitored animal welfare throughout trials. SP wrote the first draft manuscript. SP, ALW, AW and PW edited the manuscript.

## Acknowledgements

The authors would like to greatly thank the University of Liverpool IVES technical team for their support for this work in particular Mrs Karen Ryan for her help in producing the large amount of media required for bacteriology for these trials. We also thank Ceva Animal Health for their kind gift of CevaGel packs. The work in this study was supported by the UK Taxpayer via funding from the Biotechnology and Biological Sciences Research Council (BB/R008914/1).

