## Supplemental Figure 1 for "Timing and Delivery route effects of Cecal Microbiome transplants on *Salmonella* Typhimurium infections in Chickens"

### Supplemental material

#### Methods

##### *Salmonella* Typhimurium invasion of chicken epithelial cells

The chicken epithelial cell line 8E11 was used for cell invasion studies. Briefly, cells were recovered from storage in liquid nitrogen by gentle thawing at 37 $^{o}$C for 10-15 min. Then, 5-10 ml of Dulbecco's Modified Eagles Medium (DMEM) supplemented with 10% v/v foetal calf serum, 1% v/v each of non-essential amino acids, L-glutamine and Penicillin/Streptomycin (Sigma Aldrich) was added to the cells. They were then incubated for 24 h AT 37 $^{o}$C in a 5% $CO_2$ incubator. Medium was replaced every 3-4 days unti 90% confluence at which point cells were split into fresh flasks or seeded at 1 x $10^5$ cells per well in a 24 well plate. Once seeded, cells were incubated as above for 2-3 days until fully confluent. Then 24 h prior to infection cells were washed three times in PBS and fresh antibiotic free DMEM (above) was overlaid. *Salmonella* was also prepared 24 h prior to infection by picking a colony into sterile LB broth and incubating with shaking at 37 $^{o}$C. On the day of infection, cells were pre-incubated for 1 h with either 100 ul of filtered CMT or 100 ul of either Acetate or Propionate between the concentrations of 25 ug/ml to 100 ug/ml. Next, *Salmonella* was adjusted to an $OD_{600}$ of 0.1 and then 100 ul of inoculum culture was added to each test well. There were 4 wells per plate which were used as uninfected controls. Cells were incubated with *Salmonella* for 1 h after which they were washed with PBS twice and then incubated for a further 1 h with DMEM supplemented with 100 uq gentamicin to remove any planktonic and non-invaded bacteria. Cells were washed again and the lysed by the addition of 1% v/v Triton X100 for 5 min. Lysed cells were then serially diluted 10-fold and plated onto solid LB agar. Plates were incubated for 24 h and colony forming units of invaded cells were enumerated using the Miles Misra method of enumeration.

#### Result

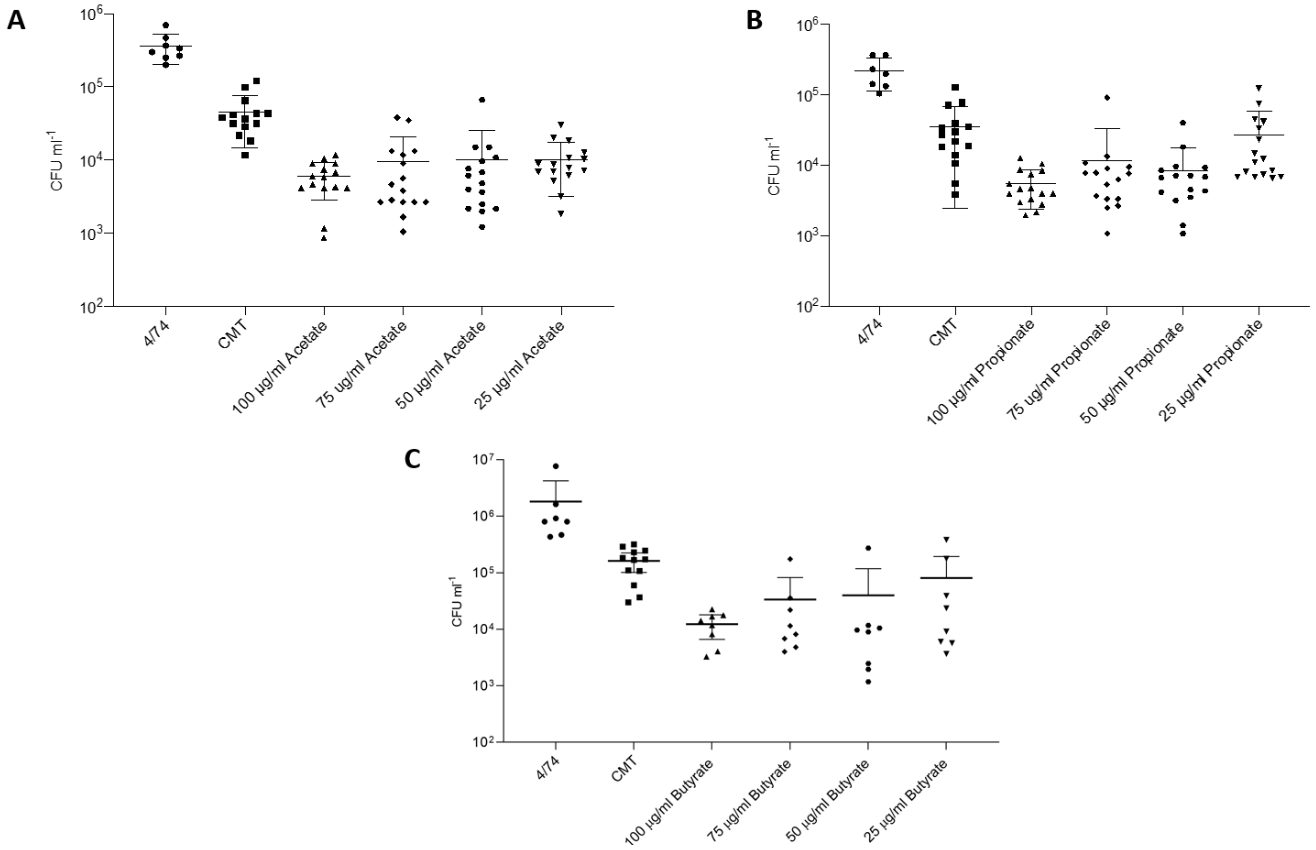


**Supplemental Figure 1: Invasion of chicken epithelial cells with *S.* Typhimurium 4/74 in the presence of CMT or Short Chain Fatty acids**. When chicken epithelial cells (8E11 cell line) were pre-incubated with either CMT or the SCFAs Acetate (A), Propionate (B) or Butyrate (C) there is a 10 -100 fold decrease in invasion of cells with ST4/74 compared to invasion controls. There is little difference in the reduced invasion of cells at varying concentrations of individual SCFA. Individual SCFAs appear to have a slightly greater effect on protection of epithelial cells than CMT alone. This could be due to other bacterial and host components within the CMT filtrate itself. For A and B graphs are representative of 4 individual experiments. For Butyrate (C) only 2 individual experiments are shown.
